# Absence of enterotypes in the human gut microbiomes reanalyzed with non-linear dimensionality reduction methods

**DOI:** 10.1101/2021.11.04.467087

**Authors:** Ivan Bulygin, Vladislav Shatov, Anton Rykachevsky, Arseny Rayko, Alexander Bernstein, Evgeny Burnaev, Mikhail S. Gelfand

## Abstract

Enterotypes of the human gut microbiome have been proposed to be a powerful prognostic tool to evaluate the correlation between lifestyle, nutrition, and disease. However, the number of enterotypes suggested in the literature ranged from two to four. The growth of available metagenome data and the use of exact, non-linear methods of data analysis challenges the very concept of clusters in the multidimensional space of bacterial microbiomes. Using several published human gut microbiome datasets, we demonstrate the presence of a lower-dimensional structure in the microbiome space, with high-dimensional data concentrated near a low-dimensional non-linear submanifold, but the absence of distinct and stable clusters that could represent enterotypes. This observation is robust with regard to diverse combinations of dimensionality reduction techniques and clustering algorithms.

## 1 Introduction

The human gut is populated by a diverse community of microorganisms. The microbiome of an individual gut settles in several years after birth and, by rough estimates, contains more than a thousand genera of bacteria [28]. Gut microbiota forms a dynamic ecosystem whose composition tends to be constant during the life of an individual but varies between individuals and may significantly depend on external and internal factors. Recently, much effort has been applied to correlate the results of deep sequencing of gut microbiota with diet, hereditary diseases, antibiotic consumption, etc. The initial sequencing of gut biota revealed that its composition tends to form discrete groups (enterotypes) consisting predominantly of taxa Bacteroides, Prevotella, and Ruminococcaceae [5]. Enterotypes have been reported in [5] as “densely populated areas in a multidimensional space of community composition” which “are not as sharply delimited as, for example, human blood groups”.

We consider enterotypes as clusters in the relative taxonomic abundance space. From a geometric point of view, clusters are defined as dense areas separated by sparse regions. Clustering is a process of assigning a finite set of objects to separate groups and identifying the natural structure of the data when the relationship between objects is represented as some metric like the Euclidean distance [66]. As mentioned in [41], a major challenge is to determine the presence of such clusters through “a thorough quantitative investigation of established clustering methods and tests for microbiome data”. The published studies rely on similar approaches yet differ in the exact number of enterotypes ranging from two [48], [89], [11], [55], [39], [54], [84], [58], [85] to three [86], [19], [80], [23], [63], and four [29]. In most papers the presence of enterotypes in the microbiome data is determined by clustering of the data into K groups, with K selected by optimization of some metric of the clustering partition quality. The resulting segregation is then visualized in the projection of the first two or three principal components from the Principal Components Analysis (PCA). This approach is quite general and may involve various intermediate steps, different clustering algorithms, and metrics; however, it has certain flaws. For example, the Partitioning Around Medoids (PAM) method used in many studies may yield erroneous results for density-based clusters. Also, widely used in the studies partition quality metrics such as the Calinski–Harabasz index [9] and the Silhouette score [64] are naturally higher for convex clusters and may fail to detect density-based partition since they rely on the estimation of inter-cluster variance and cluster centers.

Another common problem is the small size of datasets in comparison to their dimensionality. The direct application of standard clustering methods for small and high-dimensional datasets, performed in most of the works, may lead to unreliable results due to the curse of dimensionality. Clustering implies the notion of dissimilarity between data samples. When the data dimensionality increases, the concepts of proximity, distance, or nearest neighbor become less qualitatively meaningful, especially for the commonly used Euclidean or Manhattan distances [1]. For example, the distance to the nearest data point approaches the distance to the farthest data point [7]. The lack of data further amplifies this problem since the data point cloud in a high-dimensional space becomes sparse, yielding unreliable estimates of the probability density due to the non-asymptotic lower bound for the regression error [43], [36]. One straight-forward way to overcome it is to reduce the data dimensionality by PCA. However, this would allow one only to find an affine subspace containing most of the data variance. It may not be sufficient to effectively decrease the data dimensionality without significant loss of information when the data lies near a non-linear low-dimensional manifold.

Inconsistency in the number of enterotypes found in different works and aggravating factors described above undermine the very notion of enterotypes. Several studies already showed the possibility of a gradient distribution and the absence of well defined clusters [37], [88], [14]. Different structures of enterotypes between males and females were claimed by [57]. In several works, the very concept of enterotypes was described as inconsistent and uncertain. Several factors such as the variation in microbial load between samples [79], the robustness of enterotypes clusters [34], and the microbiome variation during short periods of time [41] were considered limiting to the use of the enterotype concept. However, enterotyping of the human gut has been applied in clinical research. Numerous studies claimed correlations between enterotypes and diet [40], [11], [55], [67], [49], [39], [54], [87], [85], inflammatory gut diseases [81], [10], [52], [22], [13], [31], [20], mental health [46], acne [21], stool composition [76], [78], colorectal cancer [74], circulatory diseases [38], [26], [23], psoriasis [15], and infections such as AIDS [56] and influenza [61], [70]. The idea that the information about the enterotype of an individual may be a helpful biomarker not only to correct gut diseases but also to aid other medical interventions [28] relies on the assumption that enterotypes are discontinuous clusters that are stable in time at least on the short scale; this has been challenged recently [12].

Here, we look for stable and distinct clusters in large stool microbiome datasets. The stability of clusters implies that they should not depend on the dataset, data bootstrapping, and transformations that preserve the general form of the data point cloud. Meaningful clusters should not disappear if the dataset is changed in a non-essential way. Partition of the data into distinctive clusters should correspond to high values of an appropriate clustering validity index that is applicable both for convex and density-based clusters. In other words, there should be gaps around the clusters seen as space regions with a lower concentration of points, whereas clusters correspond to areas of the highest concentration. We introduce two new intermediate steps into the common pipeline for the microbiome clustering analysis: estimation of the intrinsic dimension and manifold learning. These steps allowed us to significantly reduce the data dimensionality while preserving most of the information. Using such low-dimensional representation of the data we demonstrate the absence of stable and distinct clusters in several large datasets of 16S rRNA genotyping of stool samples.

Firstly, using Principal Component Analysis (PCA) we find a medium-dimensional linear subspace retaining almost all data cloud variation so that the dataset projected on this subspace would not significantly differ from the original dataset. Removing dimensions with low variance also acts as a filter that provides a more robust clustering [6]. However, instead of limiting the dimensionality reduction process solely to PCA, we then determine the intrinsic dimension of the projected data via Maximum Likelihood Estimation (MLE) [47]. This step allows for capturing a minimal but sufficient number of coordinates representing the most important features of the dataset. Following the manifold hypothesis [24], we suppose that microbial data cloud lies near some lower-dimensional manifold embedded in the high-dimensional abundance space. The goal of non-linear manifold learning is to obtain a low-dimensional representation of the data, supposedly lying on such a manifold while preserving most of the information. This information may be expressed as similarities or dissimilarities between data points, e.g. as a matrix of pairwise distances. non-linear projections per se are not interesting, since any data cloud could perfectly lie on a 1-dimensional submanifold.

Despite this 1-dimensional submanifold manifold yield perfect alignment in terms of minimization of the distance between the original data point and its projection, it does not preserve information in terms of pairwise distances. Therefore it is important to assess the quality of embeddings provided by manifold learning algorithms. Proper embedding should preserve the local and global structure e.g. points that are close in the original space should remain close in the embedding space. Given the intrinsic dimension, we further reduce the data dimensionality, using several manifold learning algorithms described in Materials and Methods.

For each embedding, we apply different clustering methods, namely, Spectral Clustering [69], [82], PAM and HDBSCAN [50]. Thus we obtain robust results not biased by peculiarities of the methods. HDBSCAN and Spectral Clustering are useful when the structure of the clusters is arbitrarily shaped and non-convex. We use PAM as a baseline and for comparison with the related works. For each clustering method, we iterate over a set of hyperparameter combinations to find a partition that yields the best clustering validity metrics.

Clustering metrics based on the ratio of within-cluster compactness to between-cluster separation, like the Calinski–Harabasz index (Calinski and Harabasz 1974), the Silhouette score [64], and the Davies-Bouldin index [18] cannot handle properly arbitrarily shaped clusters and noise in form of low-density points scattered around dense clusters areas. Thus, as the main metric for clustering validity, we consider Density-Based Clustering Validation (DBCV) [53], which considers both density and shape properties of clusters, takes noise into account, and is appropriate for detecting density-based clusters.

We assess the clustering partition stability using Prediction Strength [75], initially proposed for estimating the number of clusters, which tells us how well the clustering partition decision boundaries calculated on a data subset generalize the data distribution.

In addition, to be considered as a stable enterotype, a cluster should contain a sufficiently large number of data points. Hence, we do not consider spurious clusters that contain less than 5% of data assuming that such clusters are outliers or artifacts of dimensionality reduction algorithms. Indeed, they are small, depend on manifold learning algorithms, and are significantly separated from the main data point cloud. Nevertheless, they significantly impact the clustering quality metrics. To distinguish such ineligible partitions, we use the Shannon Entropy [68] of the probability distribution of data points to be in a certain cluster. All these metrics are described in detail in Supplementary Notes 6.1

To identify the optimal clustering, we compare the DBCV, Prediction Strength, and Entropy for each partition corresponding to different clustering hyperparameters. The balanced clustering partition that corresponds to a separation of the data cloud into distinct and stable clusters should produce a salient local maximum of the DBCV score, Entropy value, and Prediction Strength. Also, for each clustering partition, we calculate the Davies-Bouldin index and the Silhouette score. Lower Davies-Bouldin index and higher Silhouette score correspond to a better partition, where clusters are better in terms of compactness and separation. To demonstrate the continuous nature of the stool microbial data distribution, we construct 2d and 3d coordinate projections of the data using t-distributed Stochastic Neighbor Embedding (t-SNE) [77] and Uniform Manifold Approximation and Projection (UMAP) algorithms [51]. Hence, the workflow may be summarized as follows:

1. Use the standard PCA algorithm to obtain the initial affine subspace containing most of the data variance
2. Estimate the intrinsic dimension of the projected data using the MLE algorithm [47]
3. For each manifold learning method, find a low-dimensional representation of the data with the estimated intrinsic dimension, using the method’s hyperparameters that provide the lowest error of the learnable inverse transformation
4. Compare the clustering quality of the data for different numbers of clusters and method’s hyperparameters using the Density-Based Clustering Validation index (DBCV) [53], the Davies-Bouldin Index [18], and the Silhouette score [64]
5. Assess the clustering partition stability using the Prediction Strength [75], and evaluate partition balance using the Shannon entropy of the data distribution over clusters
6. Visualize the low-dimensional representations using t-SNE [77] and UMAP [51] algorithms

## 2 Materials and Methods

As the main source of the human gut microbiome data, we used the 16S rRNA genotype data from the NIH Human Microbiome Project (HMP) (Human Microbiome Project Consortium 2012) [33], [35] and American Gut Project (AGP) [3]. These largest available datasets provide sufficient number of data points for correct estimation of the clustering partition and constructing a manifold [60]. We used 3457 HMP samples from stool and rectum body sites downloaded from https://portal.hmpdacc.org/ and 9511 samples from AGP downloaded from https://figshare.com/ as abundance matrices. All datasets were normalized by dividing the OTUs values by the total sum of abundances for a given data sample. The numbers of objects and original dimensionality *d* in the relative taxon abundance space are presented in Tab. 1. The data were analyzed at the Order, Family, and Genus taxonomic levels (O, F, and G, respectively).

**Table 1:** Size and initial dimensionality *d* of the different datasets.

| Dataset | Size | Order $d$ | Family $d$ | Genus $d$ |
| --- | --- | --- | --- | --- |
| AGP | 9511 | 53 | 95 | 180 |
| HMP | 3457 | 51 | 96 | 172 |

The data distribution is inherently sparse due to the insufficient number of samples and noisy due to the presence of possible outliers. Noise may correspond to specific patients with extremely exotic microbial communities or be caused by sample collection or data processing flaws. Moreover, the procedures may vary between laboratories, leading to considerable batch effects. Example of a dataset-specific preprocessing is a correction for microbial blooms for the AGP dataset [2]. Therefore we can not merge several individual datasets into one. Indeed, visualization using the t-SNE and UMAP dimensionality reduction algorithms illustrates that point Fig. 1. Hence, to avoid the batch effect, we considered these datasets separately. Spurious OTU that were found in less than 1% of the samples and with a standard deviation less than 0.001 were removed.

**Figure 1:**
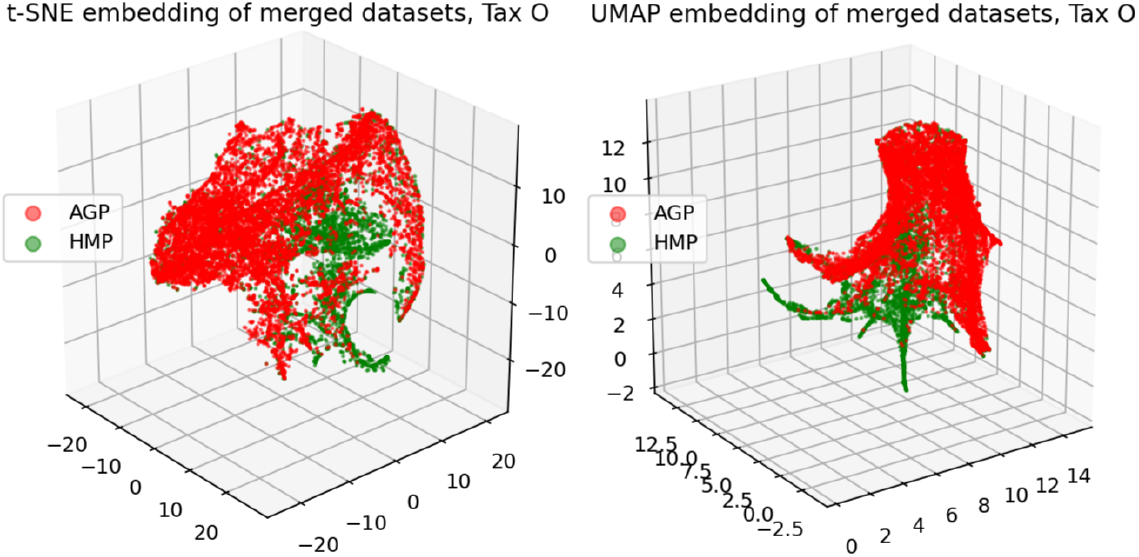
3D t-SNE and UMAP visualizations of joined datasets demonstrate batch-effect at O - Order taxonomy level (tax). Red - AGP, green - HMP dataset.

For most dimensionality reduction methods, validity indices, and metrics, implementations from the ‘scikit-learn’ library [59] were used. For the HDBSCAN clustering method and the Density-Based Clustering Validation metric the ‘hdbscan’ package [50] from the ‘scikit-learn-contrib’ was used. For the UMAP algorithm we applied implementation from the ‘umap-learn’ library [51]. Code reproducing the paper results is avaliable at https://github.com/blufzzz/Human-Gut-Microbiome-Analysis.

## 3 Results

### 3.1 Principal Component Analysis

We obtained significant dimensionality reduction with minuscule information loss by using projection on relatively many (16 through 52, dependent on the taxonomy level) principal components. The dimensionalities *d*_PCA_ of the PCA projections, are reported in Tab. 2 and defined as the number of first principal components that meet the selected threshold of 99% of the cumulative explained variance. To assess the information loss during the projection, we calculated the error of reconstruction from the projected data to the original one. The Median Absolute Error and *Q*_loc_ and *Q*_glob_ metrics (see Supplementary Notes 6.1 for details) of the reconstruction obtained using an inverse linear transformation from the projected data are presented in Tab. 3.

**Table 2:** Dimensionalities of two datasets (AGP and HMP) and three taxonomy levels (O - Order, F - Family, G - Genus). *d*_PCA_ - number of the first principal components explaining 99% variance, *d*_MLE_ - estimated intrinsic dimension.

|  | Tax O |  | Tax F |  | Tax G |  |
| --- | --- | --- | --- | --- | --- | --- |
| Dataset | $d_{PCA}$ | $d_{MLE}$ | $d_{PCA}$ | $d_{MLE}$ | $d_{PCA}$ | $d_{MLE}$ |
| AGP | 16 | 5 | 35 | 8 | 52 | 9 |
| HMP | 18 | 5 | 36 | 6 | 52 | 7 |

**Table 3:**
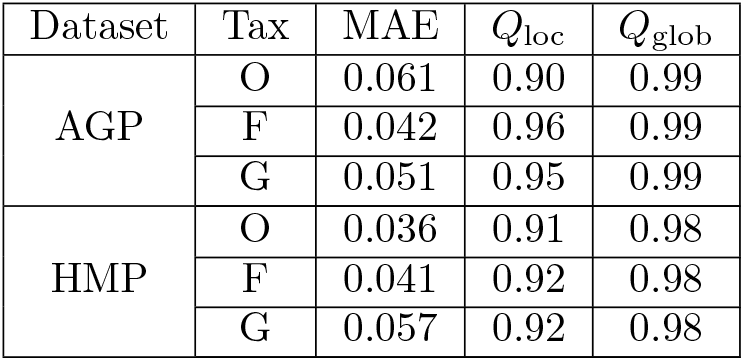
Reconstruction MAE of the linear inverse transformation from data projected on the principal components to the original space of taxon abundances and the *Q*_loc_ and *Q*_glob_ metrics (see the text for details). Notation as in Tab. 2

| Dataset | Tax | MAE | $Q_{loc}$ | $Q_{glob}$ |
| --- | --- | --- | --- | --- |
| AGP | O | 0.061 | 0.90 | 0.99 |
|  | F | 0.042 | 0.96 | 0.99 |
|  | G | 0.051 | 0.95 | 0.99 |
| HMP | O | 0.036 | 0.91 | 0.98 |
|  | F | 0.041 | 0.92 | 0.98 |
|  | G | 0.057 | 0.92 | 0.98 |

### 3.2 Estimation of the intrinsic dimension

We calculated the intrinsic dimensions *d*_MLE_ for each dataset at each taxonomic level by applying the Maximum Likelihood Estimation principle to the distances between close neighbors [47]. The intrinsic dimensions and the dimensionality after projection on principal components are presented in Tab. 2.

### 3.3 Manifold learning

For non-linear dimensionality reduction from *d*_PCA_ to *d*_MLE_, we use manifold learning algorithms, Isomap [73], Modified Locally Linear Embedding (LLE) [90], Denoising AutoEncoder (AE) [17], Spectral Embedding [69], t-distributed Stochastic Neighbor Embedding (t-SNE) [77], and Uniform Manifold Approximation and Projection (UMAP) [51]. For each manifold learning algorithm, dataset, and taxonomic level, we obtain a low-dimensional embedding. To find an optimal algorithm for a specific dataset and taxonomic level, we iterate over various combinations of manifold learning algorithm hyperparameters, for each combination assessing how well the produced embedding represents the original data. To compare the dimensionality reduction algorithms with regard to the loss of information, we construct an inverse mapping from the obtained low-dimensional manifold back to the original space using the k-Nearest Neighbours Regression [25] and estimate the reconstruction error using Leave-One-Out procedure [65] for the Median Absolute Error (MAE). This technique assesses how well points in the original space can be reconstructed given their neighbors from the embedding space.

Inverse mapping from the embedding to the original space is usually performed by a supervised learning algorithm that minimizes the reconstruction error. Selecting an algorithm, its hyperparameters, and different ways to train it would bring ambiguity into our method. Moreover, the MAE is intractable metric which does not exhibit, what aspect of data cloud has been misrepresented. Therefore we apply two additional quality criteria of dimensionality reduction [45]. These criteria, *Q*_loc_ and *Q*_glob_, represent the preservation of the “local” and the “global” structure, respectively and described in Supplementary Notes 6.1. It should be noted that *Q*_glob_ is a more important metric for the studied problem since local distortion of the data should not significantly impact the clustering partition that may be implicitly present in the data. Anyhow, for each hyperparameter combination of a dimensionality-reduction algorithm, we assess MAE, *Q*_loc_, and *Q*_glob_. Given hyperparameters that deliver the lowest MAE, we discard 10% of data points with the highest reconstruction MAE, which serves as denoising for a more stable clustering in the embedding space. It does not affect the clustering partition results, since these data points fall in the outliers category provided by the Local Outlier Factor algorithm [8]. The latter allows for detecting outliers by deviation of their local density with respect to their neighbors, from which it follows they are not related to any cluster as a densely populated area in a multidimensional space. *Q*_loc_, and *Q*_glob_ for all manifold learning methods, applied to all datasets at different taxonomic levels, are shown in Fig. 2. The MAE values are listed in Tab. 4. One may see from Fig. 2 and Tab. 4, higher taxonomy levels yield more intricate data representation with higher MAE and lower *Q*_loc_, *Q*_glob_. This is due to data becoming more sparse and dissipated in the high-dimensional space, which hinders data representation by fitting a non-linear manifold.

**Figure 2:**
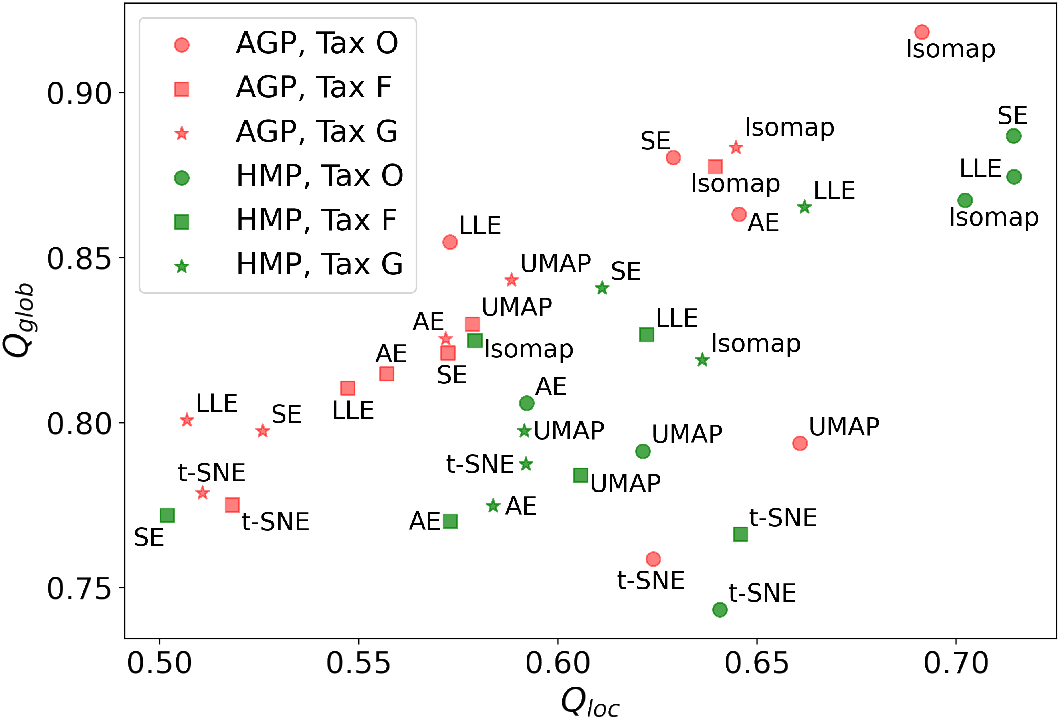
Horizontal axis - *Q*_loc_ metric (local information preservation), vertical axis - *Q*_glob_ metric (global information preservation) of the non-linear dimensionality reduction methods. Datasets (AGP and HMP) and taxonomy levels (O - Order, F - Family, G - Genus) are shown in the inset. SE - Spectral Embedding, LLE - Locally Linear Embedding, AE - AutoEncoder.

**Table 4:** The reconstruction MAE, assessed using Leave-One-Out procedure. The reconstruction is done by the independent K-Nearest Neighbour Regression of the original space of relative taxon abundances data coordinates from the non-linear embedding. Notation as in Tab. 2

| Dataset | Method | Tax O | Tax F | Tax G |
| --- | --- | --- | --- | --- |
| AGP | AutoEncoder | 0.05 | 0.19 | 0.21 |
|  | t-SNE | 0.05 | 0.19 | 0.22 |
|  | UMAP | 0.06 | 0.21 | 0.24 |
|  | Isomap | 0.06 | 0.22 | 0.25 |
|  | LLE | 0.06 | 0.22 | 0.24 |
|  | Spectral | 0.06 | 0.21 | 0.24 |
| HMP | AutoEncoder | 0.09 | 0.24 | 0.22 |
|  | t-SNE | 0.09 | 0.23 | 0.21 |
|  | UMAP | 0.11 | 0.27 | 0.25 |
|  | Isomap | 0.11 | 0.29 | 0.26 |
|  | LLE | 0.11 | 0.28 | 0.26 |
|  | Spectral | 0.12 | 0.34 | 0.28 |

### 3.4 Clustering

As described in the Introduction, we applied several clustering methods (Spectral Clustering [82], Partition Around Medoids, HDBSCAN [50]) for each embedding provided by dimensionality reduction algorithms described in Results 3.3. We use only the Euclidean metric for the manifold learning algorithms. For comparison with related works on the identification of enterotypes, we applied clustering to the original data in the high-dimensional space of taxonomic abundances with different distance metrics: Jensen-Shannon, Manhattan, Euclidean, and Bray-Curtis as in [44].

These results are included in Fig. 4 and Fig. 3. To distinguish between the presence and absence of clusters in the data we consider the following thresholds. For the Prediction Strength, we consider a score of 0.8 for moderate support as suggested in [44] and [75]. We consider all positive values of the DBCV metric. For the Silhouette score, we consider a score of 0.5 for moderate clustering as suggested in [85], [44], [27], [4]. As a threshold value for the Davies-Bouldin index, we used 0.6 for moderate clustering [18]. Under the assumption that our data capture all microbiome variations that may be possibly related to enterotypes, we do not consider small clusters that contain less than 5% of the data as natural clusters related to enterotypes.

**Figure 3:**
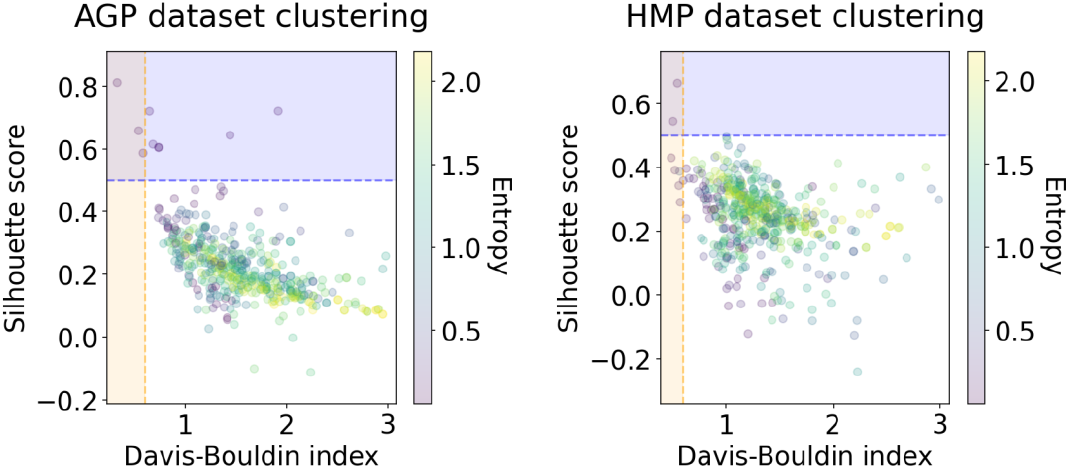
Clustering results for the AGP and HMP datasets including all three taxonomy levels (O - Order, F - Family, G - Genus) - with the corresponding Silhouette score and Davies-Bouldin index. The point color represents the Entropy of the respective partition.

**Figure 4:**
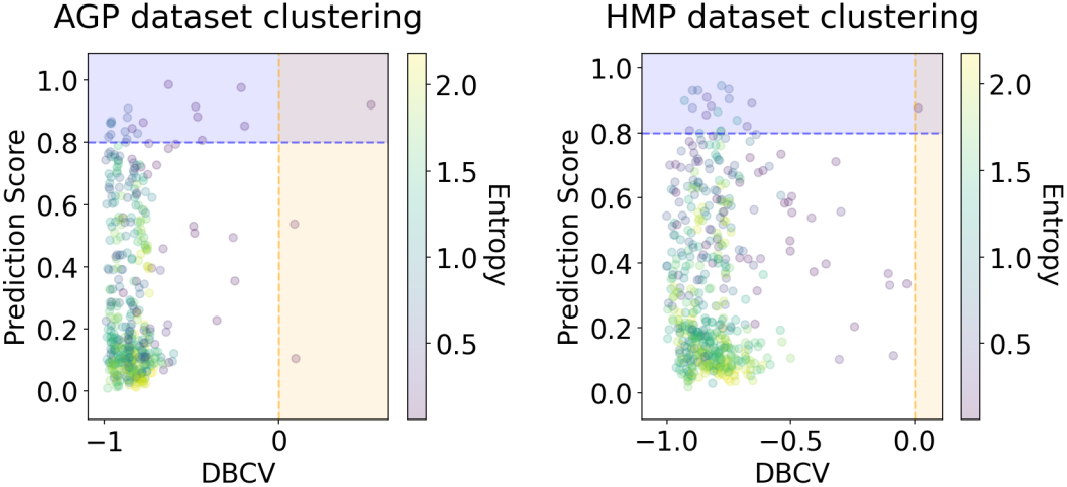
Clustering results for the AGP and HMP datasets with the corresponding DBCV index and the Prediction Strength. Notation as in Fig. 3.

To validate the ability of our methods to provide accurate clustering of data that lies in a low-dimensional non-linear manifold, embedded in a high dimensional space, we used several synthetic datasets embedded in high-dimensional space (see Supplementary Notes 6.2 for details). Since the clustering results directly depend on the algorithm hyperparameters, for each clustering method we have iterated over relevant sets of hyperparameter values. At that, for the Spectral Clustering algorithm, we considered the range from 2 to 10 as the possible number of clusters in the data, and 3, 5, and 15 as sizes of the neighborhood for computing the affinity matrix. For the PAM algorithm, we used the same range of possible clusters. For the HDBSCAN hyperparameter, we considered 10, 25, 50 as the minimal cluster size. We did not set large values for the minimal cluster size to avoid conservative clustering when more points will be considered as noise, and clusters will be restricted to more dense areas. If there is more than one partition into the same number of clusters provided by HDBSCAN, we selected the one that yields a higher DBCV index. The results of clustering demonstrated the absence of stable and distinct clusters in the data (Fig. 3 and Fig. 4).

In Fig. 3 and Fig. 4, we show the distribution of metrics over all clustering results, calculated for each taxonomic level, manifold learning method, clustering algorithm, and its hyperparameters. Clustering partitions with moderate or strong support for both metrics correspond to points lying at the intersection of the blue and orange areas at Fig. 3 and Fig. 4. All partitions that correspond to points in this area were found to consist of two highly imbalanced clusters when more than 95% of the data points are concentrated in one of them. These partitions are also inconsistent in terms of clustering validity metrics when either DBCV and Prediction Score or Davies-Bouldin and Silhouette pairs pass the thresholds but not all of them. These small clusters are unstable among different partitions of the same dataset - the mean intersection between them is lower that 30%, which asserts for high dependence on manifold learning and clustering algorithms.

Three partition that satisfies the Davies-Bouldin and Silhouette score thresholds for the AGP dataset were found in Fig. 3. One partition originates from the Locally Linear Embedding of the Order taxonomic level, where two clusters were found by the Spectral clustering algorithm. For this partition, the Davies-Bouldin index is 0.54 and the Silhouette score is 0.65, which is inconsistent with low value −0.89 of DBCV, and 0.31 for the Prediction Score. The entropy of the data mass distribution over two found clusters is equal to 0.095 which corresponds to a highly imbalanced partition where 98% of data are concentrated in one cluster.

Another two partitions of the AGP dataset into two clusters were found in the original data in high-dimensional spaces of normalized taxon-relative abundances in Order level, endowed with Jensen-Shannon and Euclidean metrics. For Jensen-Shannon the Davies-Bouldin index is 0.32, the Silhouette score is 0.81, the DBCV is −0.81 and the Prediction Score is 0.25. The entropy of the data mass distribution over two found clusters is equal to 0.099. For Euclidean distance the Davies-Bouldin index is 0.58, the Silhouette score is 0.58, the DBCV is −0.69 and the Prediction Score is 0.72. The entropy is 0.098. Another partition with DBCV and the Prediction Score above the threshold was found in the UMAP embedding of the Genus level by the HDBSCAN algorithm, presented in Fig. 4, lying at the intersection of the threshold areas. The partition consists of two clusters with the Davies-Bouldin index of 0.99, the Silhouette score of 0.26 which is incoherent with high DBCV of 0.52 and the Prediction Score of 0.92. Entropy is equal to 0.06 which corresponds to an imbalanced partition where 98% of data belong to one cluster.

For the HMP dataset, three highly imbalanced partitions into two clusters that are satisfying the metrics thresholds were found. The first one in Fig. 3 was generated by the HDBSCAN clustering algorithm in the Spectral embedding at the Family taxonomic level. This partition yields the Davies-Bouldin index of 0.54, the Silhouette score of 0.66, DBCV of −0.64, the Prediction Score of 0.21, and the Entropy of 0.091. The second one in Fig. 3 was found in the Locally Linear embedding of the Order level by the Spectral Clustering algorithm with the Davies-Bouldin index of 0.50, the Silhouette score of 0.54, DBCV of −0.84, the Prediction Score of 0.45, and the Entropy of 0.1. The only clustering partition that satisfies the Prediction Score and DBCV thresholds for the HMP dataset at Fig. 4 was found in the t-SNE embedding at the Order level, by the HDBSCAN algorithm. For this clustering, the Davies-Bouldin index is 1.01, the Silhouette score is 0.11, DBCV 0.01, the Prediction Score is 0.87, and the Entropy is 0.05, which indicates 99% of data concentrated in one cluster.

Together, these results imply that various clustering methods along with different manifold learning algorithms yield highly imbalanced partitions. We do not consider clusters that contain less than 5% of the data as enterotypes. We attribute these small clusters to artifacts of manifold learning algorithms since there are evidences [16], [42] that common dimensionality reduction techniques may fail to faithfully represent the original point cloud distribution, introducing substantial distortion into the data. We observe that these small clusters are not stable, depending on the clustering method and the manifold learning algorithm. Hence, the stool metagenomes can hardly be divided into stable and distinct clusters that could be referred to as enterotypes. Our simulation on a synthetic dataset in Supplementary Notes 6.2, reaffirms that distinct and stable clusters related to enterotypes were not found because of their absence in the data rather than methodology flaws.

Despite the lack of distinct and stable clusters in the data, we demonstrate that human gut microbial communities vary continuously along a low-dimensional manifold. We observe the structure of such a manifold by mapping the point cloud, projected on the principal components, on a 2-dimensional plane, using t-SNE in Fig. 5 and UMAP in Fig. 6.

**Figure 5:**
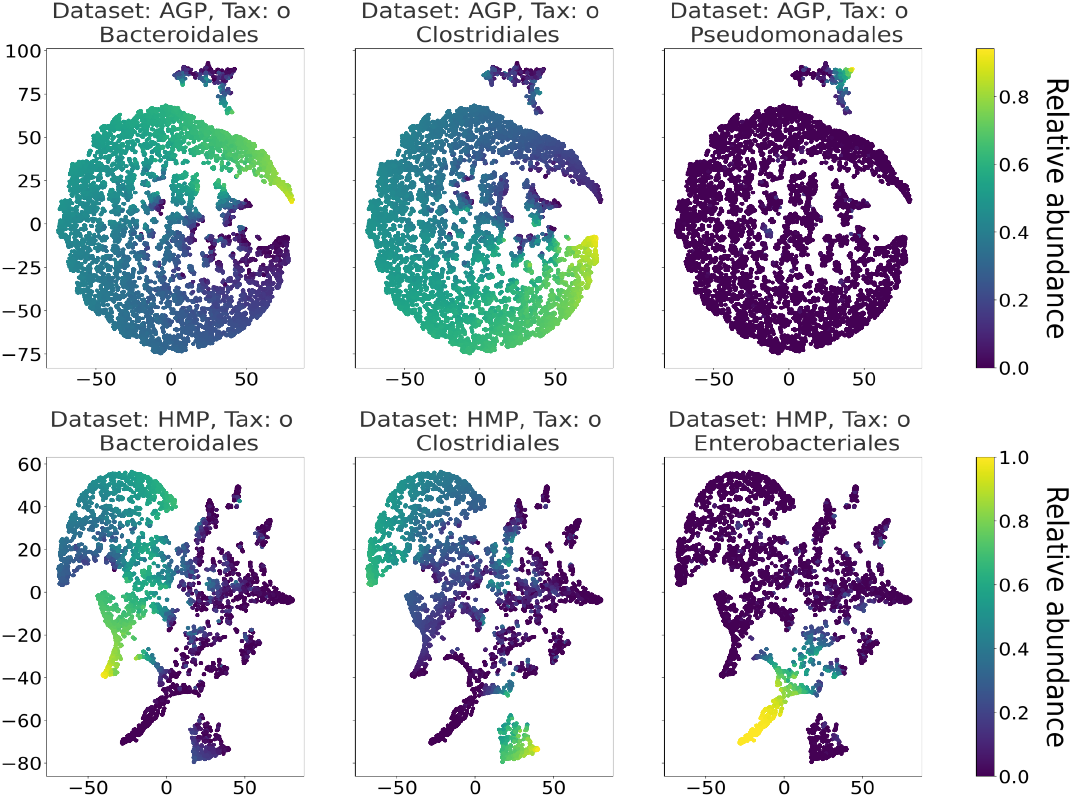
2D t-SNE visualization of AGP and HMP datasets for the Order taxonomy level. Colors reflect the relative abundance of specific taxa, see the headers.

**Figure 6:**
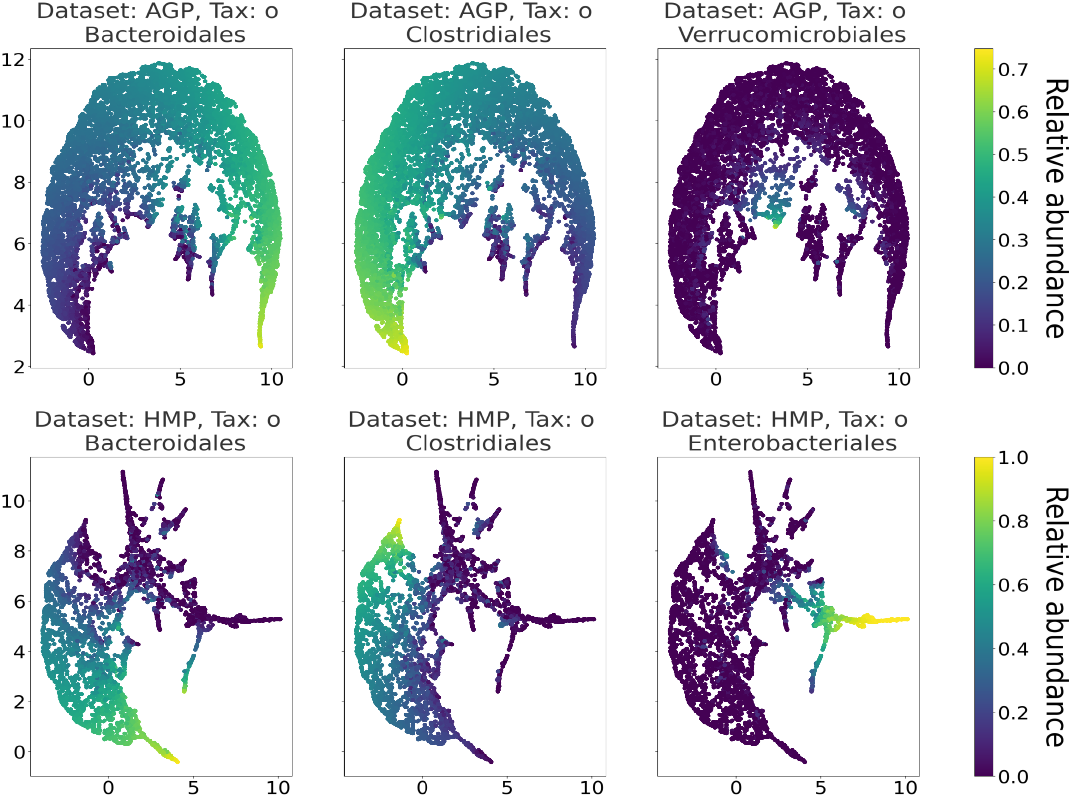
2D UMAP visualization of AGP and HMP datasets for the Order taxonomy level. Colors reflect the relative abundance of specific taxa, see the headers.

To demonstrate the continuous specificity of the data points distribution, we colored points corresponding to specific taxon relative abundances. Salient parts of the manifold correspond to higher concentrations of specific OTUs. Small clusters observed in Fig. 5 and Fig. 6 are not related to enterotypes being the direct result of specific methods hyperparameters, that may lead to tearing off the salient part of data manifold.

## 4 Discussion

Our results support that the metagenome distribution is continuous rather than discrete and lies on a low-dimensional non-linear manifold embedded in the original high-dimensional space of relative taxon abundances. We posit that most of the previous observations may have been artifacts caused by limitations of linear methods that have been applied for the analysis of non-linear and high-dimensional microbial data. Also, overfitting in the data data points. Small sizes of datasets lead to unstable clustering, especially if it is performed in a high-dimensional space. One may suggest an intuitive explanation of why positive clustering results were widespread in previous works. In most of them, small datasets were used, which makes the total number of abnormal microbial patterns negligible. We suppose that datasets demonstrating moderate clustering in related works have been sampled from high-density areas in the general taxonomic abundance space. Such areas correspond to prevalent types of microbial patterns, whereas areas with low density correspond to abnormal and unusual ones. Still, successful attempts to correct the human gut microbiota were made, e.g. fecal microbiota transplantation to treat Clostridium difficile infection (CDI) [71], inflammatory bowel disease (IBD) [72], and obesity [83]. Connecting the distribution of microbiome abundances and features of the microbial manifold with lifestyle, nutrition, or disease may be a promising direction for further research.

## 6 Supplementary Notes

### 6.1 Metrics

The Silhouette score *S_i_* [64] for a data point *i* is defined as:

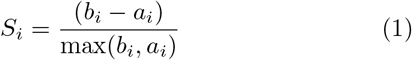

In Eq. 1, *a_i_* is the mean intra-cluster distance and *b_i_* is the mean distance between the sample point *i* and the data points from the nearest cluster *C_k_* that sample *i* is not a part of.

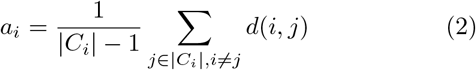

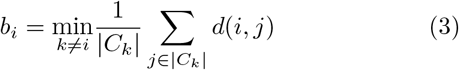

Here in Eq. 2, 3, |*C_i_*| is the number of points in a cluster *C_i_* and *d*(*i*, *j*) is the distance between data points *i* and *j*. The total Silhouette score is measured as the mean of all *S_i_* and bounded between −1 for incorrect clustering and +1 for dense and well-separated clusters. The Davies-Bouldin index DB [18] in Eq. 4 measures the average similarity of the distance between clusters with the size of the clusters themselves. It estimates the cohesion based on the distance from the points in a cluster to its centroid and the separation based on the distance between centroids. Values closer to zero indicate better partitions.

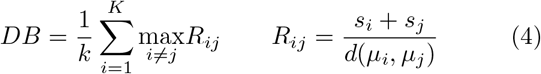

In Eq. 4, *s_i_* is the average distance between each point of cluster *i* and a centroid *μ_i_* of that cluster, *K* is the total number of clusters, and *d*(*μ_i_, μ_j_*) is the distance between centroids *μ_i_* and *μ_j_* of the clusters *i* and *j*.

Density-Based Clustering Validation was proposed as a metric for assessing the clustering partition quality for non-convex clusters. The metric is based on the Hartigan model of Density Contour Trees [30] and provides values between −1 and +1, with greater values indicating a better density-based clustering solution. See [53] for details.

To assess disbalance in a clustering partition, we use the Shannon Entropy [68] *E* of the distribution that indicates the probability *p_i_* of one of *n* data points to fall into a cluster *i* with |*C_i_*| data points in it, see Eq. 5. Higher *E* values indicate more balanced partition.

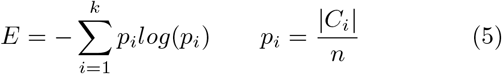

We estimate the reconstruction error of inverse mapping 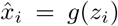 from the manifold learning embedding *z* ∈ *Z* to the original space of relative taxon abundances as the normalized Median Absolute Error (Eq. 6):

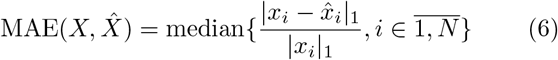

Selecting and supervised training of the inverse mapping algorithm *g* brings ambiguity into the method. To mitigate it, we use additional scale-independent rank-based quality criteria for dimensionality reduction proposed in [45]. Given a distance metric *d*(*i*, *j*), two rank matrices are constructed 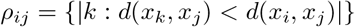 for the original data points *x_i_* ∈ *X* and 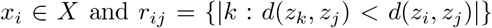 for their low-dimensional embeddings *z_i_* ∈ *Z*. Then, a co-ranking matrix *Q* is built, where 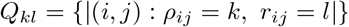 indicates the total number of instances when the *k*-th neighbor for some point became *l*-th. Thus, all non-diagonal elements of this matrix correspond to changes in arrangement of the embedded points compared to the original data. The co-ranking matrix contains all information on how the data structure is distorted in a low-dimensional representation. This information is represented further as two scalars *Q*_loc_ and *Q*_glob_ that range from 0 (bad) to 1 (good) and reflect the preservation of the “local” and “global” structure of the data cloud. See [45] for details.

For similarity measure between the predicted clustering partition *X* and given ground-truth labels *Y* we utilize Adjusted Rand Index [32], based on the Rand Index (RI) [62]. Given the predicted 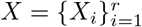 and the true 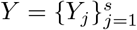 clustering partitions into *r* and *s* clusters respectively, the Rand Index is non-zero for random independent partitions *X* and *Y* and 1 for the same partitions up to labeling permutations. The Adjusted Rand Index is “adjusted” version of RI such its values are close to 0 for random labeling. In Tab. 7 a contingency table is presented, where *n*_*ij*_ denotes the common number of points in clusters *X_i_* and 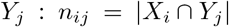. Given *a_i_*, *b_j_*, *n_ij_* from Tab. 7, the Adjusted Rand Index (ARI) is defined as Eq. 8. The 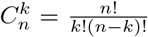 denotes a *k*-combination from a set of *n* elements.

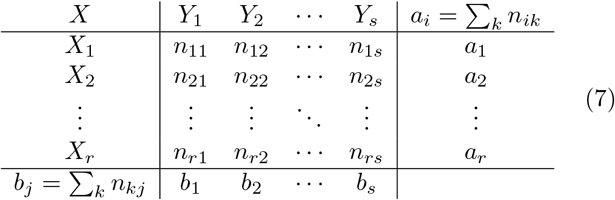

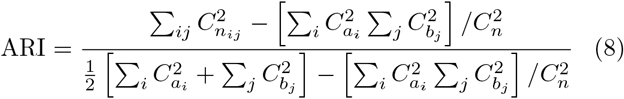

### 6.2 Synthetic dataset

In this section, we provide the results of our method on synthetic data. The positive control datasets with clusters were created similarly to [44]. We generated nine synthetic datasets with the number of clusters *k* ranging from two to four, with different dimensionality *d* equal to the O,F and G taxonomy levels (53,96 and 180 correspondingly). Each dataset consist of 3000 data points that represent vectors of un-normalized abundances of mock OTUs. 90% of data points were sampled from one of *k* multivariate gaussian distributions, each reflecting a cluster, and other 10% are sampled from a distribution with large variance, representing noise. To compare estimated clustering partitions with the ground-truth labels, we used the Adjusted Rand Index [32] (see Supplementary Notes 6.1 for details). The dimensionality that retains 99% of the cumulative explained variance after the PCA projection, estimated intrinsic dimension, the Median Absolute Error and *Q*_loc_, *Q*_glob_ metrics are reported in Tab. 5.

**Table 5:** Original dimensionality *d*, dimensionality *d*_PCA_ after PCA projection and estimated intrinsic dimensionality *d*_MLE_ of synthetic datasets. For each dataset MAE, *Q*_loc_ and *Q*_glob_ metrics were computed for the PCA projection with respect to original data.

| $d$ | $k$ | $d_{\text{PCA}}$ | $d_{\text{MLE}}$ | MAE | $Q_{\text{loc}}$ | $Q_{\text{glob}}$ |
| --- | --- | --- | --- | --- | --- | --- |
| 53 | 2 | 2 | 2 | 0.045 | 0.89 | 0.98 |
| 53 | 3 | 2 | 2 | 0.074 | 0.85 | 0.98 |
| 53 | 4 | 4 | 3 | 0.068 | 0.86 | 0.98 |
| 96 | 2 | 2 | 2 | 0.080 | 0.85 | 0.97 |
| 96 | 3 | 6 | 4 | 0.086 | 0.86 | 0.97 |
| 96 | 4 | 22 | 8 | 0.082 | 0.87 | 0.98 |
| 180 | 2 | 109 | 21 | 0.091 | 0.86 | 0.97 |
| 180 | 3 | 58 | 13 | 0.086 | 0.86 | 0.98 |
| 180 | 4 | 21 | 7 | 0.095 | 0.85 | 0.98 |

In Fig. 7 we display the clustering metrics distribution over all partitions that have been calculated for each dataset with a given number of clusters *k* and dimensionality *d*, each manifold learning method, and each clustering algorithm with a different combination of its hyperparameters. Clustering partitions with moderate or strong support for both metrics correspond to points lying at the intersection of the blue and orange areas in Fig. 7. For each dataset the most accurate clustering partition was found in the UMAP embedding by the HDBSCAN clustering algorithm. Such partitions have the Adjusted Rand Index and the Prediction score equal to 1, the Davies-Bouldin index lower than 0.3, the Silhouette score higher than 0.5 and DBCV higher than 0.8. Together, these metrics assert presence of stable and distinct clusters in the data for every number of clusters *k* and dimensionality *d*.

**Figure 7:**
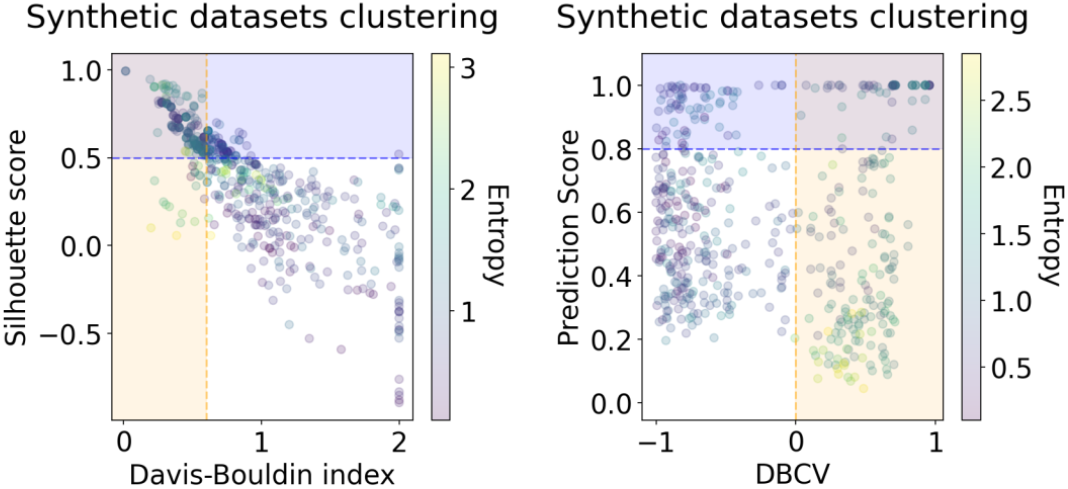
Silhouette score and Davies-Bouldin index (left), DBCV index and Prediction Strength (right) of clustering partitions for all nine synthetic datasets with the number of clusters *k* equal to 2, 3 and 4, and the dimensionality *d* of O, F and G taxonomy levels, equal to 53, 96 and 180.

